# miRNA858b Inhibits Proanthocyanidin Accumulation by Repression of *DkMYB19* and *DkMYB20* in Persimmon

**DOI:** 10.1101/783431

**Authors:** Sichao Yang, Meng Zhang, Liqing Xu, Zhengrong Luo, Qinglin Zhang

## Abstract

Persimmon proanthocyanidin (PA) biosynthetic had been reported to be regulated by several transcription factors, but the miRNAs function involved in this process was poorly understood. We identified a miRNA858b that putatively targeted two R2R3-MYB transcription factors, *DkMYB19*/*DkMYB20*. Transcript accumulation of *DkMYB19*/*DkMYB20* and miRNA858b showed contrasting divergent expression patterns during fruit development. DkMYB19/DkMYB20 were confirmed to be localized in the nucleus. The interaction between miRNA858b and *DkMYB19*/*DkMYB20* were experimentally validated by 5’ RNA ligase-mediated RACE and LUC enzyme activity detection. Overexpression of miRNA858b led to the down-regulation of *DkMYB19*/*DkMYB20* which reduced the accumulation of PA, whereas the reduced miRNA858b activity that up-regulated the *DkMYB19*/*DkMYB20* resulted in high levels of PA in STTM858b transient expression in leaves *in vivo*. Similarly, the transient transformation of miRNA858b in fruit wafers *in vitro* also reduced the accumulation of PA by repressing the *DkMYB19*/*DkMYB20*, while the up-regulation of *DkMYB19*/*DkMYB20* enhanced the accumulation of PA in STTM858b or *DkMYB19*/*DkMYB20* transient transformation in fruit wafers. PA content decreased after overexpression of miRNA858b in *Arabidopsis* wild type and *DkMYB19*/*DkMYB20* in persimmon leaf callus consisted with the above results. These findings suggested that miRNA858b repressed the expression of *DkMYB19*/*DkMYB20* which contribute to PA accumulation in persimmon.

## Introduction

Persimmon (*Diospyros kaki*), one of East Asia’s major fruit crops, accumulates large amounts of PA in its fruit “tannin cells” that cause strong astringency sensation in fresh fruits. Persimmon varieties are classified into two groups according to genetic characteristic of natural de-astringency: pollination constant nonastringent (PCNA) persimmons, which is a nonastringent (NA) mutant phenotype that loses its astringency naturally on the tree during fruit development, so that fruits can be edible at the firm stage. Non-PCNA type need artificial treatment to remove astringency with carbon dioxide gas or ethanol vapor or by drying after peeling before consumption. Both the allelotype of astringent (A) and NA are controlled by a single gene known as the *AST*/*ast* allele (Akagi *et al*., 2009b, 2011).

In Japanese PCNA type persimmon, the growth of PA in cells gradually terminates during the early developmental stages of persimmon fruit, with the loss of astringency principally occurring via PA dilution as the fruit grows larger (“dilution effect”). Astringency removal in the Chinese PCNA persimmon fruit presented the characteristics of both the Japanese PCNA and non-PCNA types: the process is related to both “dilution effect” and “coagulation effect” (the conversion of soluble tannins into insoluble tannins) (Yonemori *et al*., 2003). Understanding the mechanism of astringency removal will contribute to molecular breeding of novel PCNA persimmon cultivar. Recently, several key genes and MYB transcriptional factors involved in astringency removal in Chinese PCNA persimmon had been identified. Mo *et al*. (2016a) showed that *DkADH* and *DkPDC* genes specific to the natural astringency-loss trait of Chinese PCNA persimmon fruit. Min *et al*. (2012) showed that two hypoxia-responsive *ERF9*/*ERF10* were involved in separately regulating the *DkPDC2* and *DkADH1* promoters in regulating persimmon de-astringency. *DkMYB6* was identificated as a putative transcriptional activator, induced by high CO2, which is involved in persimmon fruit deastringency, by operating on both *DkPDC* structural genes and *DkERF* transcription factors (Fang *et al*., 2016). Meanwhile, pyruvate kinase genes (*DkPK*) are involved in the natural loss of astringency in Chinese PCNA persimmon via enhancement of the transcript levels of the *DkPDC* and *DkADH* genes (Guan *et al*., 2016; 2017). Furthermore, Xu *et al*. (2017) revealed that *DkALDH2a* and *DkALDH2b* negatively regulate the natural deastringency in Chinese PCNA.

Proanthocyanidin (PA, also called condensed tannin) biosynthesis involves synthetic pathways controlled by both structural genes and transcription factors (TFs). Those structural pathway genes encoding enzymes concerned anthocyanin pathway including 3-deoxy-Darabino-heptulosonate 7-phosphate synthase (*DAHPS*), 3-dehydroquinate synthase (*DHQS*), 3-dehydroquinate dehydratase/shikimate 5-dehydrogenase (*DHD/SDH*), shikimate kinase (*SK*), 5-enolpyruvylshikimate 3-phosphte synthase (*EPSPS*), chorismate synthase (*CS*), phenylalanine ammonia lyase (*PAL*), cinnamate-4-hydroxymate (*C4H*), 4-coumarate: coenzyme A ligase (*4CL*), chalcone isomerase (*CHI*), chalcone synthase (*CHS*), dihydroflavonol 4-reductase (*DFR*), flavanone 3-hydroxylase (*F3H*), flavanone 3’-hydroxylase (*F3’H*), flavonoid 3’5’ hydroxylase (*F3’5’H*) and two important genes as anthocyanidin reductase (*ANR*) and leucoanthocanidin reductase (*LAR*) for PA-specific pathway (Akagi *et al*., 2009b; Akagi *et al*., 2011; Akagi *et al*., 2012). Other genes involved in PA accumulation, such as *TT12* encoding a multidrug and toxic compound extrusion family transporter (Debeaujon *et al*., 2001), *TT19* encoding a glutathione S-transferase (GST; Kitamura *et al*., 2004), *Auto-Inhibited* H+ -*ATPase Isoform* (*AHA10*) encoding a proton pump involved in vacuolar biogenesis (Baxter *et al*., 2005), a membrane-localised multi-drug and toxic compound extrusion (MATE) transporter targets epicatechin 3’-*O*-glucoside in *Medicago truncatula* (Zhao and Dixon, 2009), laccase-like polyphenol oxidase (*TT10*) for PA polymerization (Zhao *et al*., 2010) have been identified.

Furthermore, some transcription factors such as the Myb-like protein, which regulates transcription of the structural genes involved in PA biosynthesis, have been identified in a few plant species. In *Arabidopsis*, *TT2*, a Myb transcription factor that controls the transcription of *ANR*, *DFR*, *Auto*-*Inhibited* H+ -*ATPase isoform* (*AHA10*), and *TT12* have been characterized (Lepiniec *et al*., 2006). In the grape berry, two MYB activators such as *MYBA7* and *MYBA1* act as inducers of both anthocyanins and flavonols (Matus *et al*., 2008, 2017), *VvMYBF1* induce flavonols in grape tissues by upregulating the transcription level of the flavonol synthase (*FLS*) gene (Czemmel *et al*., 2009). Another two transcription factors *MYBPA1* and *MYBPA2* control the expression of *ANR* and *LAR* that involved in the accumulation of precursors of flavonoids concerned flavonols, PA, and anthocyanins (Czemmel *et al*., 2012). *MYB114* acts as a key regulator mediating the production of flavonols by converting dihydroxy flavonols to flavonols (Tirumalai *et al*., 2019). The *VvMYBC2-L1* and *VvMYB5b* have been reported to negative regulate PA accumulation (Huang *et al*., 2014) and contribute to the regulation of anthocyanin and PA biosynthesis in developing grape berries respectively (Deluc *et al*., 2008). In coleus, a MYB transcription factor named *SsMYB3* is involved in the regulation of PA biosynthesis (Zhu *et al*., 2015). In apple and poplar, *MdMYB9* and *PtMYB134* presented the ability to activate *ANR* promoters and to partner with heterologous bHLH co-factors from those two plants (Gesell *et al*., 2014). In *Medicago truncatula*, *MtPAR* is an MYB family transcription factor that functions as a key regulator of PA biosynthesis (Verdier *et al*., 2012).

In persimmon, two MYB transcription factors, *DkMYB2* and *DkMYB4*, have been reported to involve in PA regulation in persimmon by activating the *DkF3’5’H*/*DkANR* and *DkANR*/*DkLAR* transcription, respectively (Akagi *et al*., 2009b, 2010a). Subsequently, a *DkbZIP5* could directly regulate *DkMYB4* in an ABA-dependent manner through recognizing ABA-responsive elements in the promoter region of *DkMYB4* (Akagi *et al*., 2012) and a *DkMYC1* might be an important bHLH transcription factor for the regulating of PA biosynthesis (Su *et al*., 2012; Naval *et al*., 2016; Nishiyama *et al*., 2018) have been verificated. Meanwhile, DkMYB2 or DkMYB4 could interact with DkMYC1 and DkWDR1 to compose a MBW ternary complexes which involved in regulating PA accumulation in fruit (Naval *et al*., 2016).

MicroRNAs (miRNAs) are important elements of post-transcriptional gene regulation in eukaryotes. In plant, miRNAs are processed from a primary miRNA transcript (pri-miRNA), which includes a foldback structure, by the nuclear RNase DICER-LIKE 1 (DCL1) and its accessory proteins SERRATE (SE) and HYPONASTIC LEAVES1 (HYL1) (Achkar *et al*., 2016). In addition to transcription factors, miRNAs play crucial roles in the regulatory networks by modulating gene expression at the posttranscriptional level (Voinnet, 2009). A search of literatures evidence suggests that the majority of plant miRNAs are known to target transcription factors so as to play critical regulatory roles in multiple biological processes, such as the development, primary and secondary metabolism, as well as stress responses (Sharma *et al*., 2016; Wang *et al*., 2016; Wu, 2013). Particularly, some miRNA involved in regulating anthocyanin, flavonoid and PA biosynthesis have been reported, such as the miRNA156-*SPL9* network directly influences anthocyanin production through targeting genes encoding *PAP1* and dihydroflavonol 4-reductase (Gou *et al*., 2011; Cui *et al*., 2014), miRNA408, a positive regulator of photomorphogenesis, leads to increase anthocyanin accumulation in *Arabidopsis* seedings (Zhang *et al*., 2011, 2014), and miRNA828 or TAS-siRNA 81(-) negatively regulates the anthocyanin biosynthesis (Hsieh *et al*., 2009; Luo *et al*., 2012; Yang *et al*., 2013), In *Halostachys capsica*, miRNA6194 targets the flavanone 3b-hydroxylase mRNA (*F3H*) (Yang *et al*., 2015), In *Rauwolfia serpentina*, the miRNA396b and miRNA828a target the mRNAs coding for anthocyanin regulatory C1 protein (Prakash *et al*., 2016).

At present, it was well known that several MYBs from persimmon fruit were targets of miRNAs, but it was not known how exactly such targeting might influence changes in the anthocyanin, flavonol and PA levels. The PA is recognized as a important secondary metabolism in the persimmon during the fruit development stages, the function of miRNA in PA accumulation process is far to known. Here, we screened a miRNA858b from the miRNA database of persimmon, which was predicted to target two MYB genes *DkMYB19*/*DkMYB20*. MiRNA858b and *DkMYB19*/*DkMYB20* molecular dynamic expression were performed together with PA content changes in Chinese PCNA persimmon. Our results suggested that miRNA858b and its target gene *DkMYB19*/*DkMYB20* showed a divergent expression pattern in the fruits of persimmon during development. In addition, our findings provided evidence to that miRNA858b-MYB played an important role in the regulation of PA biosynthesis in persimmon.

## Materials and Methods

### Materials

Chinese PCNA persimmon genotype ‘Eshi 1’ (*D. kaki* Thunb.; 2n = 6x = 90) was planted in the Persimmon Repository, Huazhong Agricultural University, Wuhan, China. Fruit flesh samples from uniform fruits free of visible defects were collected at 2.5, 5, 10, 15, 20, and 25 weeks after bloom (WAB) (Fig. 1). Twelve fruits per sample type were collected for each following treatment, which repeated three times. All samples were frozen immediately in liquid nitrogen and then stored at **-**80 °C after sampling. Tissue culture seedlings of ‘Gongcheng Shuishi’ (PCA genotype) were grown on MS (1/2N) media in a growth chamber with a 12-h photoperiod and 24 °C temperature. The tobacco (*Nicotiana benthamiana*) plants were grown under 16 h light/8 h dark cycles at 21 °C.

**Fig. 1.**
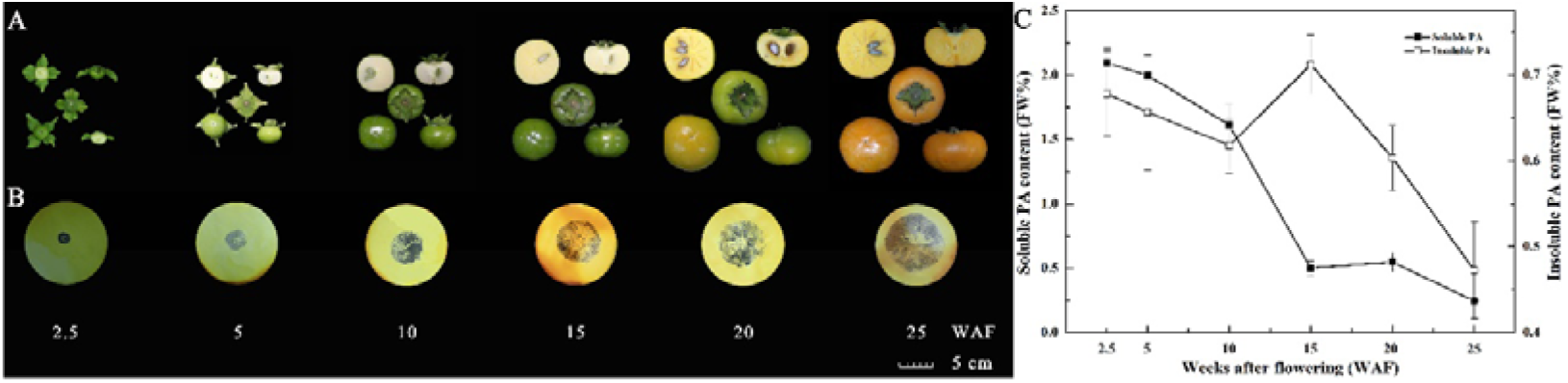
Measurement of the PA content of persimmon fruits at different developmental stages. A. Representative photos showing sampled fruits at six stages, 2.5, 5, 10, 15, 20 and 25 WAB. B. Analysis of soluble PA content in the persimmon fruits based on an imprinting method. C. Quantitative measurement of soluble and insoluble PA in the fruits using Folin-Ciocalteau method.

### PA content determination

To measure the tannin content, approximately 0.1 g of flesh from each of three fruit was homogenized with 80% (V/V) MeOH in a 2 ml tubes. The homogenate was centrifuged at 5000 g for 10 min and pellet was washed again with the same solvent. The combined supernatant, containing soluble tannins, was made up to 1 ml with further solvent. The pellet containing insoluble tannins was then suspended in 1% (V/V) hydrochloric acid in methanol (1% HCl-MeOH) and 65 °C water bath for 1 hour. The extract was centrifuged and the pellet was washed again with 1% HCl-MeOH. The combined supernatant, i.e. resolubilized tannin fractions, was adjusted to 1 ml with solvent. Tannin concentration of both soluble and resolubilized fractions were measured by the Folin-Ciocalteau method (Oshida *et al*., 1996) and was calculated as (+)-catechin equivalents. In brief, a suitable aliquot of each fraction was made up 8.5 ml with water, and mixed with phenol reagent (0.5 ml) and saturated sodium carbonate solution (1.0 ml). Then, after 1 h, the adsorbence was determined at 725 nm using as a blank containing 8.5 ml water and reagents only by a UV-2450 spectrophotometer (Shimadzu, Japan). The PA content was also examined by staining tissues with DMACA (p-Dimethylaminocinnamaldehyde) solution (0.6% DMACA and 1% 6 mol/L HCl in methanol) (Li *et al*., 1996). The infiltrated fruit wafers were decolorized in 30% acetic acid in ethanol for 12-20 h. Next, they were washed with 75% ethanol and then stained blue in DMACA solution for 2 min. The deeper blue means higher PA content.

### RNA extraction and cDNA synthesis

Total RNA was isolated from fruit flesh using RNAPlant Plus Reagent (Tiangen Biotech Co., Beijing, China). The RNA quality and quantity were assessed by gel electrophoresis and Nanodrop 2000 spectrophotometry (Thermo Fisher, USA). Three biological replicates were performed for each sample. For gene isolation, first-strand cDNA was generated using 2.0 μg total sample RNA with EasyScript One-Step gDNA Removal and cDNA Synthesis SuperMix Kit (TransGen, Beijing, China) according to the manufacturer’s protocol. For gene expression analysis, cDNA was synthesized from 1.0 μg of each RNA sample using PrimeScript RT Kit with gDNA Eraser (TaKaRa, Dalian, China).

### Prediction of target and miRNA precursor secondary structure

The target gene prediction was performed by psRNATarget program (http://plantgrn.noble.org/psRNATarget/) based on the miRNA database (accession number: SRP050516, Luo *et al*., 2015) and transcriptome database (Chen *et al*., 2017). The main principle focus on the degree of sequence homology between the miRNAs and targets. RNA Folding Form software was used to construct the miRNA precursors secondary structure (Zuker, 2003).

### Isolation and cloning of the *DkMYB19*/*DkMYB20* genes

In the transcriptome database of Chinese PCNA persimmon established in our laboratory (Chen *et al*., 2017), two unigenes (CL4549.Contig2_All, Unigene36868_All) were predicted and screened out as candidate target genes of miRNA858b, the full-length sequences of the *DkMYB19* and *DkMYB20* genes were isolated by RACE based on the original unigene sequences. The primer sequences used for RACE and cloning are described in Table S1.

### Sequence alignment and phylogenetic analysis

The gene sequences were translated with online software (http://linux1.softberry.com/). Deduced amino acid sequences of homologous genes with 41 other species were retrieved from National Center for Biotechnology Information (NCBI). A phylogenetic tree was constructed using the neighbor-joining (NJ) method with the MEGA5 software (Tamura *et al*., 2011).

### Quantitative Reverse Transcription PCR (qRT-PCR)

Quantitative reverse transcription PCR was performed with a LightCycler R^®^ 480 II System (Roche Diagnostics, Switzerland). The PCR reaction mixture (10 μl total volume) included 5 μl SYBR Premix Ex Taq II (TaKaRa, Dalian, China), 3.5 μl ddH_2_O, 1.0 μl diluted cDNA, and 0.25 μl of each primer (10 μM). The PCR conditions were as follows: 5 min at 95 °C; 45 cycles of 95 °C for 5 s, 58 °C for 10 s, and 72 °C for 15 s; and a melting temperature cycle with constant fluorescence data acquisition from 65 to 95 °C. Each sample was assayed in quadruplicate, and *DkActin* (accession no. AB473616) was used as the internal control (Table S1).

### RNA ligase-mediated 5’ RACE

RNA ligase-mediated rapid amplification of 5’ cDNA ends (RLM-5’ RACE) was conducted with a GeneRacer kit (Invitrogen, Carlsbad, CA, USA) to map the cleavage sites of target transcripts. Three replicates of l g total RNA, isolated from pooled samples of persimmon fruits, were used for ligating 5’ RACE RNA adaptors at 15 °C overnight. Gene-specific primers (Table S1) were designed to conduct 5’ RACE PCR. The PCR products were cloned into the pEASY-Blunt Simple vector (TransGen Biotech, China) for sequencing.

### Validation of the target gene of miRNA858b

To verify whether *DkMYB19* and *DkMYB20* could be cleaved by pri-miRNA858b, we conduct a co-transformed into *nicotiana benthamiana* leaves using the *Agrobacterium*-mediated (GV3101) transfection system. Vector pGreenc 0800-Luc, which contains a reporter gene LUC, was used as the expression vector. The pri-miRNA858b sequence and the ORF sequence of *DkMYB19/DkMYB20* without the termination codon was fused with the LUC reporter gene by double enzyme method. The primers was in the Table S1. The experimental procedure was according to Lee *et al*. (2008).

### Subcellular localization of DkMYB19 and DkMYB20

The complete open reading frame (ORF) of *DkMYB19*/*DkMYB20* without the termination codon was amplified using the respective primers (Table S1) in order to conducted the fusion constructs 35S-DkMYB19::GFP, 35S-DkMYB20::GFP according to Guan *et al*. (2016) and the 35S-GFP vector was acted as a positive control. The fusion and control plasmids were transferred into *Agrobacterium tumefaciens* strain GV3101 by heat shock and then transiently transformed into leaves of 6-week-old *N. benthamiana* plants. Three days later, the fluorescence signal was detected under a fluorescence microscope (Nikon 90i, Japan) (Zhao *et al*., 2019).

### Transient expression in persimmon leaves *in vivo*

A transient over-expression system was utilized to assess the role of miRNA858b and *DkMYB19*/*DkMYB20* in regulating the PA biosynthesis of persimmon leaves *in vivo*. Full-length pri-miRNA858b, *DkMYB19* and *DkMYB20* gene sequences were inserted into the plant expression vector pMDC32 by homologous recombination according to Mo *et al*. (2016b), while the GFP inserted pMDC32 vector was used as the control, the primers were listed in Table S1. The short tandem target mimic (STTM) of miRNA858b (STTM858b) module was inserted in pMDC32 vector between the 2×35S promoter and the 35S terminator using a reverse PCR procedure according to Tang *et al*. (2012) with primers described in Table S1. The construct was transferred into persimmon leaves via a previously described *Agrobacterium*-mediated method, with minor modifications (Mo *et al*., 2015). The *Agrobacterium* cells collected after centrifugation and then were re-suspended to an optical density (OD) at 600 nm of 0.75. The transformed leaves still grew on the tree before they were isolated for qRT-PCR and PA content analysis. Eight days later after injection of those constructs into leaves, 100 mg tissue from each infiltrated leaf was collected for further determination, with a total of ten single leaf replicates.

### Transient transformation in persimmon fruit wafers *in vitro*

We introduced the transient transformation system in persimmon fruit wafers. Wafers of 1 cm diameter and 0.5 cm thickness were incubated for 1 h with *Agrobacterium* infecting solution carrying constructs. The wafers were then transferred to flter papers wetted by MS liquid medium in tissue-culture dishes, and placed in an incubator at 25 °C for 3 d. All of the treatments including all genes and the empty vector were performed with three biological replicates. At each sampling point, the waferss were dried on flter papers, frozen in liquid nitrogen and then stored at −80 °C for further utilization (Zhu *et al*., 2018).

### Transformation in *Arabidopsis* wild type

The *Agrobacterium tumifaciens* GV3101 line containing 2×35S::pri-miRNA858b was introduced into the *Arabidopsis thaliana* wild type according to the planta transformation procedure of Zhang *et al*. (2006). Seeds from the T1 generation transgenic plants were grown and selected on MS medium containing kanamycin (50 mg/L).

### Genetic transformation in persimmon leaf callus

Transformations of persimmon were performed with *Agrobacterium tumifaciens* according to the methods reported by Tao *et al*. (1997). Pieces of leaf were infected with the transformed *A. tumefaciens*, and callus tissue was regenerated on MS(1/2N) solid medium containing 50 mg/L kanamycin as a selective antibiotic.

### Yeast two-hybrid assays

To detect proteins that may interact with *DkMYB19*/*DkMYB20* in persimmon, yeast two hybrid (Y2H) assays were performed. The full-length coding sequence of *DkMYB19*/*DkMYB20* were amplified by PCR using cDNAs as the template (Table S1), and then were ligated into the pGBKT7 (binding domain, BD) bait vectors via the *Bam*H c and *Sma* c double digestion. For the yeast-two-hybrid screening, an expressed CPCNA ‘Luotian-Tianshi’ fruit cDNA library was recombined into the pADGAL4 prey vector. The resulting bait and prey vectors were transformed into yeast strains AH109 by the LiAc/DNA/PEG method according to the Matchmaker® Gold Yeast Two-Hybrid System User Manual, respectively. Co-transformed yeast cells were firstly plated on the SD medium lacking leucine and tryptophan (SD-Leu-Trp), and incubated at 30 ^◦^C for 3-5 days. Transformed yeast cells containing pGBKT7-53 + pGADT7-T, pGBKT7-Lam + pGADT7-T, pGBKT7 constructs were used as a positive, a negative and a blank control, respectively. The pGBKT7-*DkMYB19*/*DkMYB20* were transformed in yeast cells as a self-activation activity assay. Single colonies were patched on quadruple-selection SD medium lacking adenine, histidine, leucine, and tryptophan (SD-Ade-His-Leu-Trp), followed by incubation at 30 ^◦^C for 3–5 days. Yeast colony PCR using 5′ and 3′ PCR primers (Supplementary Table 1), were performed to determine the presence of inserts in the prey, pGADT7-Rec clones.

Plasmids were isolated from yeast colonies derived from the QDO selective media using the Dr. GenTLE® (from Yeast) High Recovery, and the “prey” vectors containing inserts of candidate interactors, were isolated by transforming into Trans1-Blue Chemically Competent Cell and plating on LB with ampicillin (Amp), (selective for only pGADT7-Rec clones). Colonies were picked, cultured in LB/Amp (overnight) and the plasmids were purified. The protein-protein interactions (PPIs) were confirmed by co-transforming Y2HGold with the “bait” (*DkMYB19* in pGBKT7) clone together with the interactor “prey” clone (in pGADT7-Rec) and plated on QDO. To check for any false positive interactions, the empty “bait” vector was cotransformed with the interactor prey clone and plated as above. The pGADT7-Rec clones were sequenced in the forward and reverse directions using T7 and 3′AD primers (Eurofins, India). The sequences of the identified interacting protein were subjected to BLASTN and BLASTX (NCBI, http://www.ncbi.nlm.nih.gov/) analyses for identification and confirming the correct orientation of the interactor sequences and to rule out any false-positive or large ORFs in the wrong reading frame (Ramalingam *et al*., 2015).

## RESULTS

### The PA accumulation lasted to the final stage of fruit development in Chinese PCNA

The imprinting method was used to determine PA content levels in persimmon fruits. The sections were deeply stained at the beginning of fruit development (2.5 WAB, weeks after bloom), when the young fruits of ‘Eshi 1’ were small in size. With the progression of development, the fruits grew quickly and became increasingly larger until reaching their largest sizes at 25 WAB (Fig. 1A). The fruits were still darkly stained until 15 WAB, but then began to diffuse at 20 WAB. At the last experimental stage, 25 WAB, the fruits were only lightly stained (Fig. 1B). To confirm the imprinting results, quantitative measurements of soluble and insoluble PA contents in the fruits were carried out using the Folin-Ciocalteau method. The soluble PA concentration in the fruits was 21.0 mg/g at 2.5 WAB, but quickly decreased until 15 WAB (5 mg/g), with only a slight increase at 20 WAB, then decreased to the lowest level (1.72 mg/g) at 25 WAB, that accounted less than 0.2% of the fruit weight, implying the fruits at this point have already lost their astringency. Insoluble PA, which had remarkably lower levels than soluble PA, showed a trend of continuous decline except for an increased process from 10 WAB to 15 WAB, it reached its peak value (7.1 mg/g) at 15 WAB and lowest value (4.7 mg/g) at 25 WAB (Fig. 1C).

### Isolation and sequence analyses of the *DkMYB19*/*DkMYB20* in Chinese PCNA persimmon

*DkMYB19* with 633 base pairs was isolated and was predicted to encode a protein of 210 amino acid residues, while *DkMYB20* comprised of 771 base pairs and was predicted to encode a protein of 256 amino acid residues. Both proteins contained an N-terminal R2R3-DNA binding domain (Fig. S2). To determine the evolutionary relationship of *DkMYB19* and *DkMYB20* with other plant R2R3-MYB nucleic acid sequence, a neighbor-joining phylogenetic tree was assembled using the full-length nucleic acid sequences of *DkMYB19*, *DkMYB20* and all functionally tested MYBs involved in the regulation of anthocyanin and PA biosynthesis, as well as MYBs concerned in flavonoid biological processes. Our results showed that *DkMYB19* and *DkMYB20* were all clustered into the PA-related MYB clade, and *DkMYB19* was closely related to *TaMyb14* (AB252152.1), *VvMYB5b* (AY899404.1), *OsMYB4* (D88620.1), the protein sequence share 35%, 34% and 30% similarity with DkMYB19 respectively. *DkMYB20* was phylogenetically closer to *DkMYB2*, which has been previously described as a PA biosynthetic regulator (Akagi *et al*., 2010a), and the protein sequence similarity rate is 98% (Fig. 2). These results suggested that the *DkMYB19* and *DkMYB20* genes encode putative MYB proteins which may involved in regulating PA biosynthesis, in Chinese PCNA.

**Fig. 2.**
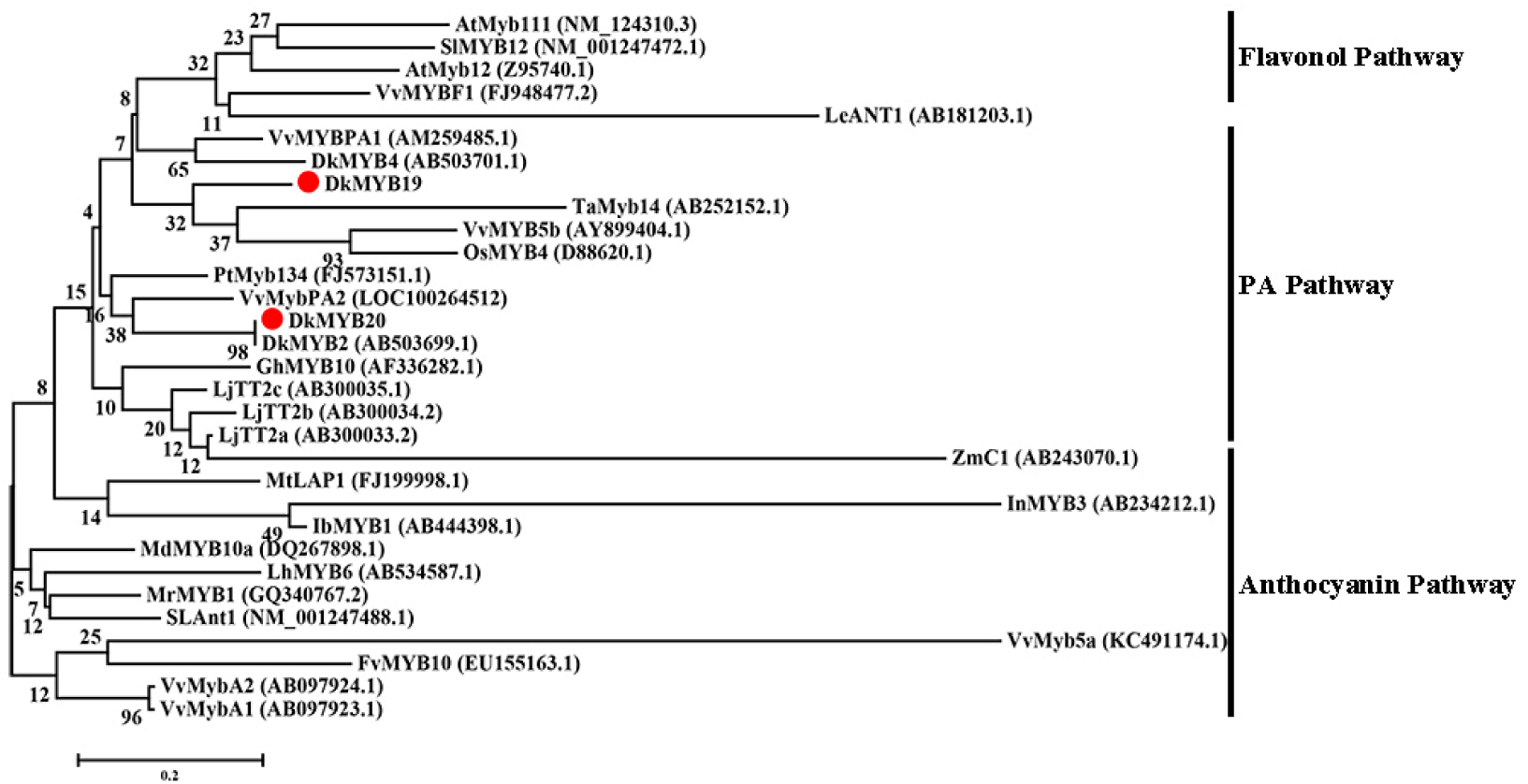
Phylogenetic tree of DkMYB19 and DkMYB20 homologs constructed with the neighbor-joining algorithm using MEGA5. Branches are labeled with the gene names from different plant species. Bootstrap values were calculated from 2000 bootstrap replicates.

### miRNA858b and *DkMYB19/DkMYB20* shared contrasting divergent expression patterns during fruit development

To gain insight into the expression patterns of the *DkMYB19*, *DkMYB20* and miRNA858b genes, their relative expression levels in developing persimmon fruit were examined using quantitative reverse transcription-polymerase chain reaction (qRT-PCR). Interestingly, the expression of *DkMYB19* and *DkMYB20* showed a divergent pattern compared with that of miRNA858b during the fruit development stage. The relative expression of miRNA858b was very low before 20 WAB, then sharply increased to the maximum value at 20 WAB, followed by a slight increase at 25 WAB (Fig. 3).

**Fig. 3.**
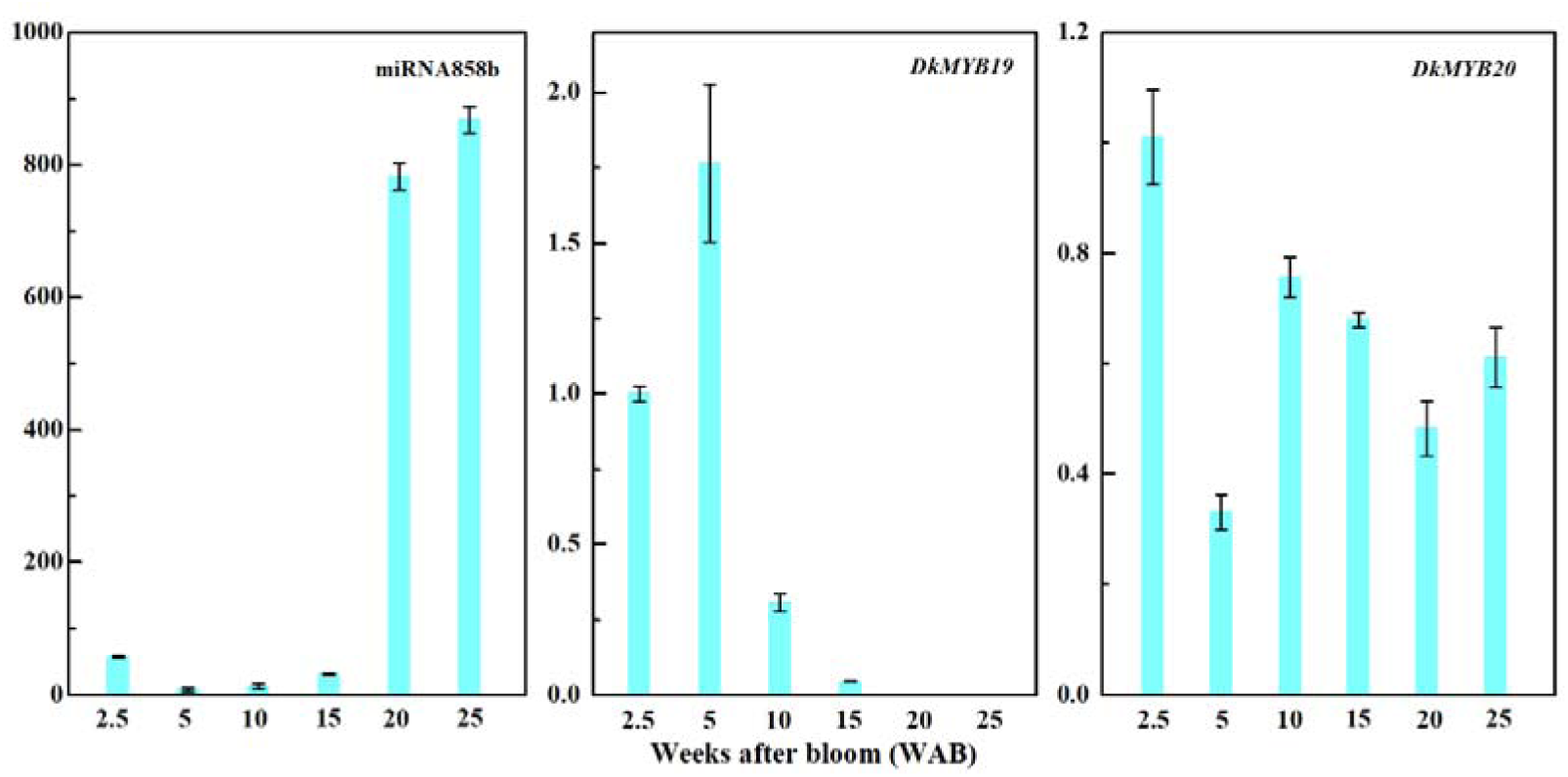
Expression of miRNA858b, *DkMYB19* and *DkMYB20* during persimmon fruit development. Error bars indicate the standard deviation (n = 3).

In addition, the transcript level of *DkMYB19* presented an up-regulation process at 5 WAB, then it tended to decrease from 5 to 25 WAB (Fig. 3). The level of *DkMYB20* decreased approximately 70% at 5 WAB, followed by a noticeable up-regulation at 10 WAB, then it showed a trend of down-regulation from 10 to 20 WAB, followed by a slight up-regulation at 25 WAB (Fig. 3).

### Validation of the target gene of miRNA858b

The PCR products of pri-miRNA858b were cloned and sequenced to confirm the incorporation of persimmon-specific miRNA and miRNA* in the backbone. From the Mfold output, the guiding strands of the miRNAs were considered without changing any other nucleotide. Predicted miRNA/miRNA* duplexes are shown in Figure 4A.

**Fig. 4.**
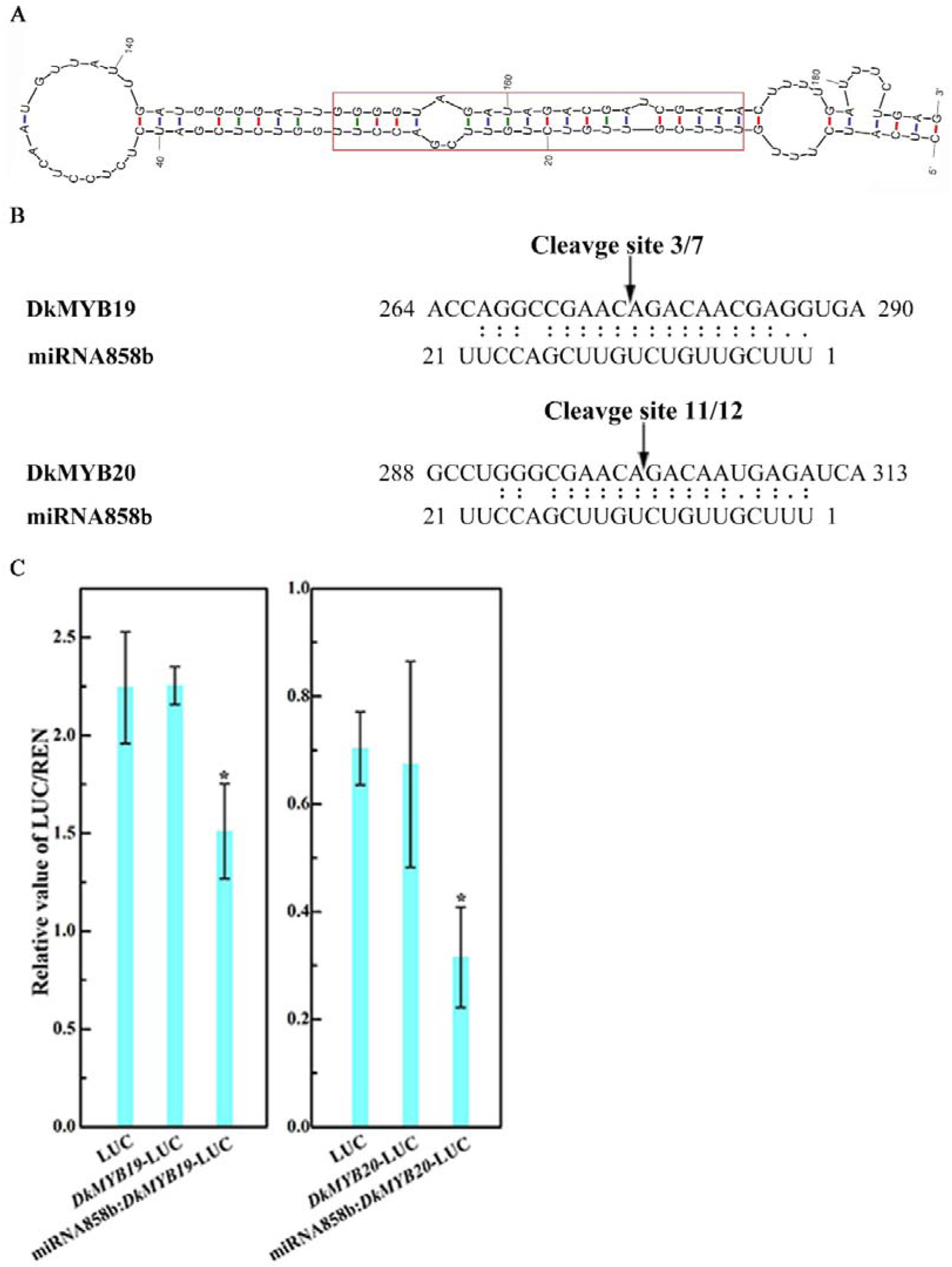
Experimental validation of predicted targets of miRNA858b. A. Secondary structure of the miRNA858b precursor predicted by Mfold. The red box represents the sequence of the miRNA/miRNA* duplex. B. Sequence comparison between the target genes, *DkMYB19* and *DkMYB20*, and their corresponding miRNAs, miRNA858b, respectively. The arrows indicate the cleavage sites, and the numbers show the frequency of clones sequenced. C. Co-transformation of *DkMYB19/DkMYB20* and miRNA858b in tobacco leaves. Recombinant vectors were transformed into tobacco leaves by *Agrobacterium* strain GV3101. Dual-LUC assay showing interaction of miRNA858b and its target gene. Relative LUC activity was normalized to the Renilla (REN) luciferase. Error bars indicate SD (n =4). (P < 0.05).

To confirm the target cleavage sites of pri-miRNA858b, the downstream sequence of the predicted target cleavage sites were used to design outer and inner primers for 5’ RACE, to obtain upstream sequences including the miRNA target region. Sequencing of the 5’ RACE PCR clones demonstrated that pri-miRNA858b has unique cleavage sites in their corresponding targeted sequences at 10-11 nt sites (Fig. 4B).

To verify the accuracy of RLM-5’ RACE result, we applied co-transformation technology in tobacco leaves using the vector pGreenc 0800-Luc containing the LUC reporter gene. The LUC/REN of leaves inoculated with GV3101-pGreenc 0800-Luc (control) was same with the leaves inoculated with GV3101-pGreenc 0800-Target-Luc, in which the target sequence was fused upstream of the LUC gene.

The LUC/REN was decreased in leaves co-transformed with GV3101-pGreenc 0800-Target-Luc and GV3101-pGreenc 0800-pri-miRNA858b. In agreement with RLM-5’ RACE sequencing, the pri-miRNA858b could cleave the candidate targets (Fig. 4C).

### DkMYB19 and DkMYB20 were localized in the nucleus

Most R2R3 MYB genes are exclusively localized and function in the nucleus (Stracke *et al*., 2001). The deduced DkMYB19/DkMYB20 protein sequences contain putative nuclear localization signals, suggesting the nuclear localization of DkMYB19/DkMYB20. To confirm this, 35S-DkMYB19::GFP, 35S-DkMYB20::GFP fusion vectors were constructed and transiently transformed into tobacco leaf cells via agroinfiltration. 35S:GFP alone, driven by the 35S promoter was used as a negative control. The subcellular localizations of the DkMYB19/DkMYB20-GFP fusion proteins were determined by visualization using a confocal microscope. Similar to other R2R3 MYB transcription factors, the DkMYB19/DkMYB20-GFP fusion protein was localized exclusively in the nucleus, whereas GFP (control) was uniformly distributed throughout the cell (Fig. 5). These results indicated that DkMYB19/DkMYB20 encode nuclear-localized proteins.

**Fig. 5.**
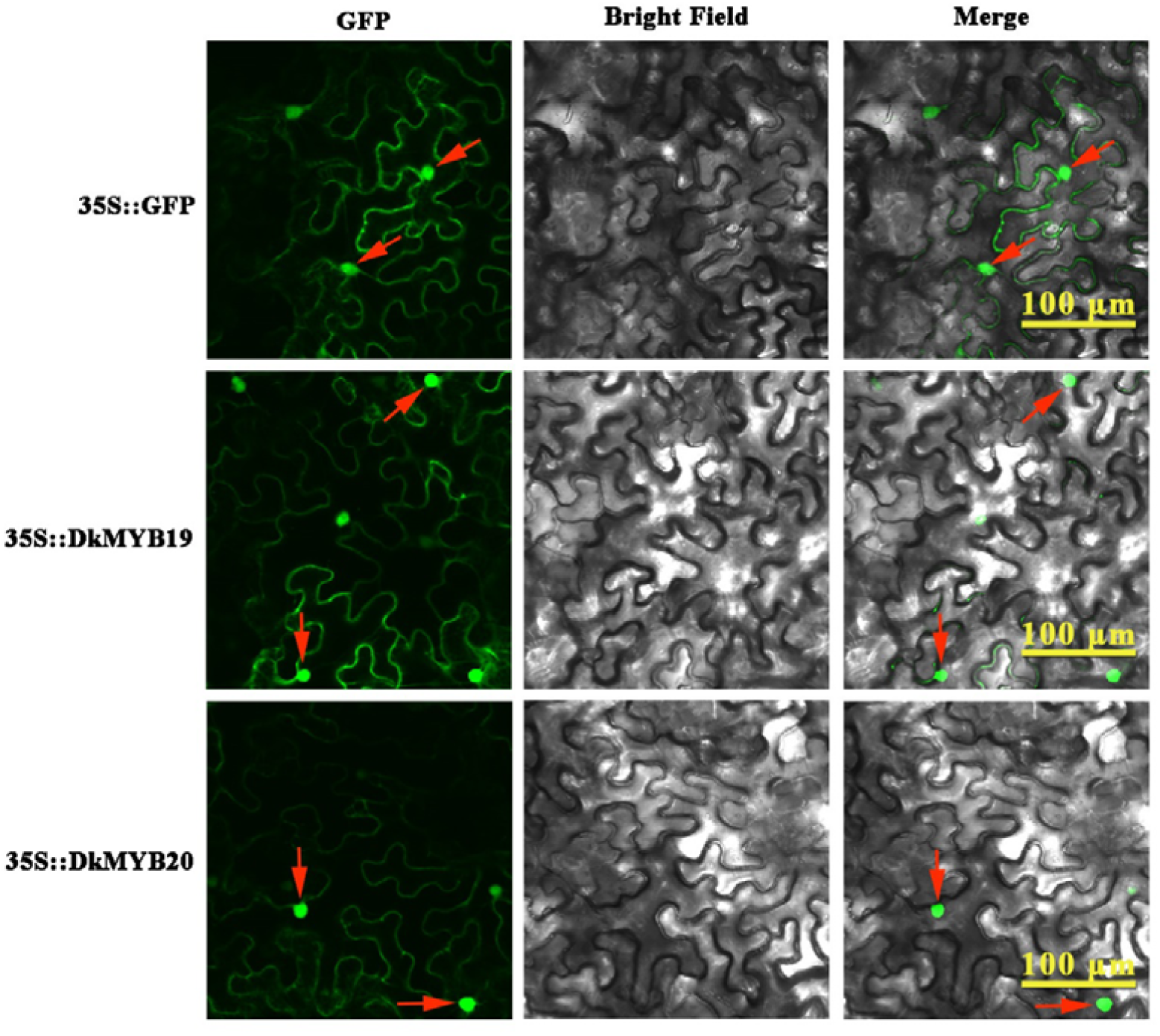
Subcellular localization of the DkMYB19/DkMYB20 proteins. DkMYB19/ DkMYB20-GFP fusion or green fluorescent protein (GFP) alone was transiently expressed in tobacco epidermal cells under the control driven by cauliflower mosaic virus (CaMV) 35S promoter and observed under a confocal microscope. Brightfield photographs were taken for cell morphology, dark field was used for green fluorescence, and in combination. Bar, 100 μm. The red arrows represents the nucleus.

### Transient expression in persimmon leaves and fruit wafers

The matches between miRNA858b and *DkMYB19*/*DkMYB20* mRNA indicate that miRNA858b is a suppressor of DkMYB19/DkMYB20 protein biosynthesis. As shown in Fig. 8, when pri-miRNA858b was infiltrated in leaves *in vivo*, the expression levels of miRNA858b showed sharply up-regulation 21.63-fold relative to the control, while the *DkMYB19*/*DkMYB20* gene expression was suppressed, which leaded to significantly down-regulation of *DkDHD/SDH*, *DkPAL*, *DkCHS*, *DkCHI* and slightly decreased of *DkF3’H*, *DkF3’5’H*, *DkDFR*, *DkANS*, *DkANR* and *DkLAR* that encode the enzymes of PA biosynthesis pathway. As expected, the levels of soluble and insoluble PA were all significantly decreased in the persimmon leaves after infiltration with pri-miRNA858b (Fig. 6A). When pri-miRNA858b was infiltrated in ‘Eshi 1’ fruit wafers, the DMACA staining exhibited distinct shallows as compared with control (Fig. 7A), which meant the PA content was decreased (Fig.7B). Morever, a miRNA858b blocking construct, STTM858b, was generated and infiltrated in leaves *in vivo*, we found the expression of miRNA858b was sharply down-regulated 88.67% against to the control, while *DkMYB19/DkMYB20* gene expression was up-regulation, together with the up-regulation of *DkDHD*/*SDH*, *DkPAL*, *DkCHS*, *DkCHI*, *DkF3’H*, *DkF3’5’H*, *DkDFR*, *DkDFR*, *DkANS*, *DkANR*, *DkLAR*. As a consequence, soluble and insoluble PA content were both significantly increased in the persimmon leaves after STTM858b infiltration (Fig. 6B). After STTM858b was infiltrated in fruit waferss, the DMACA staining exhibited distinguished darker compared with control (Fig. 7A), which also implying the PA content increasing (Fig. 7B).

**Fig. 6.**
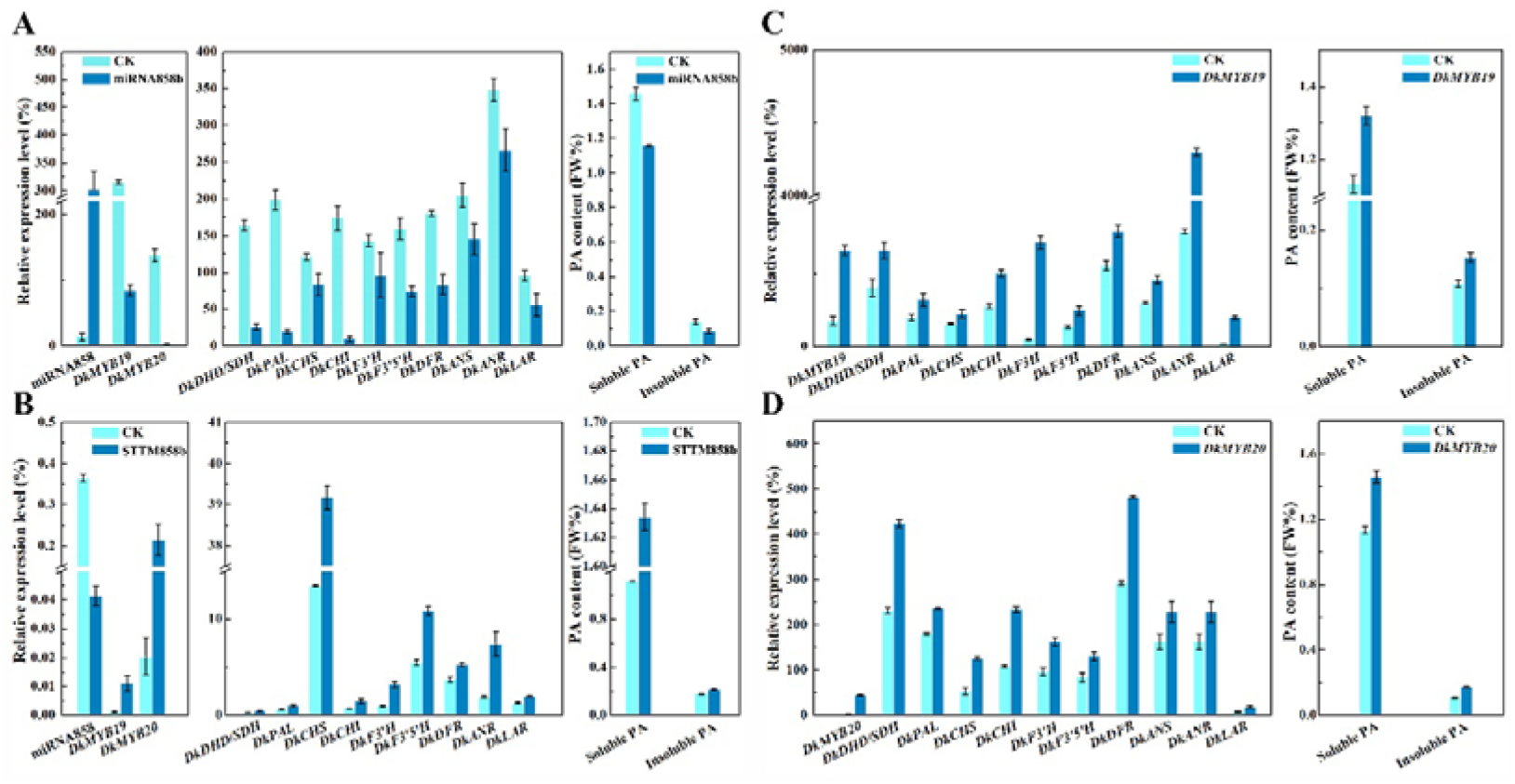
Transient expression of miRNA858b, STTM858b, *DkMYB19* and *DkMYB20* in persimmon leaves *in vivo*. A, Analysis of the transcript level of the miRNA858b, *DkMYB19*, *DkMYB20*, PA pathway genes and its PA content variation after transient over-expression of primiR858 in ‘Eshi 1’ persimmon leaves *in vivo* 10 days after agroinfiltration. B, Analysis of the transcript level of miRNA858b, *DkMYB19*, *DkMYB20*, PA pathway genes and its PA content variation after transient expression of STTM858b in ‘Eshi 1’ persimmon leaves *in vivo* 10 days after agroinfiltration. C, Analysis of the transcript level of *DkMYB19*, PA pathway genes and its PA content variation after transient expression of *DkMYB19* in ‘Eshi 1’ persimmon leaves *in vivo* 10 days after agroinfiltration. D, Analysis of the transcript level of *DkMYB20*, PA pathway genes and its PA content variation after transient expression of *DkMYB20* in ‘Eshi 1’ persimmon leaves *in vivo* 10 days after agroinfiltration.

**Fig. 7.**
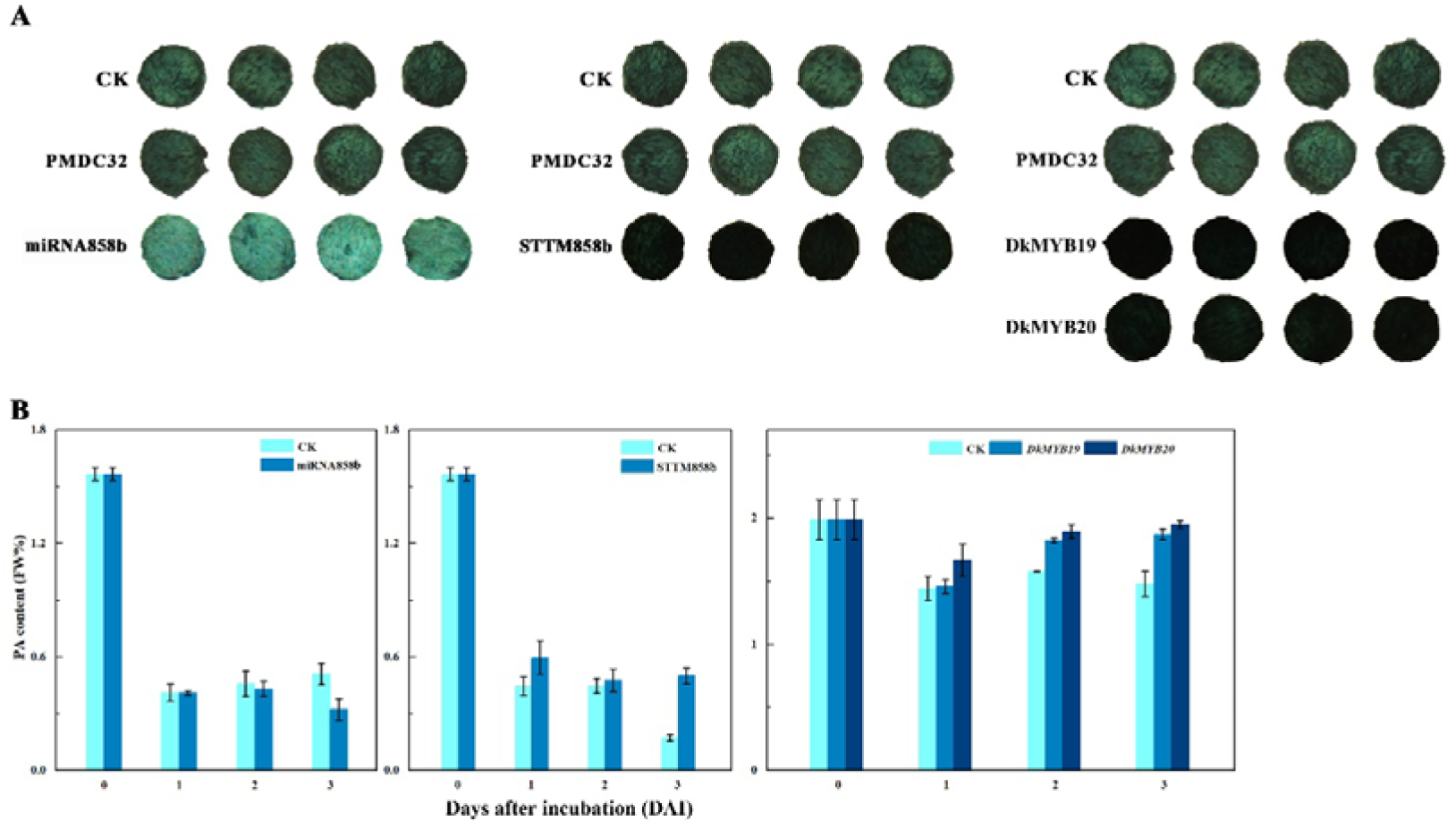
PA concentration variation in persimmon fruit wafer transformation of miRNA858b, *DkMYB19* and *DkMYB20* by DMACA staining (A) and Folin-Ciocalteau (B) method.

**Fig. 8.**
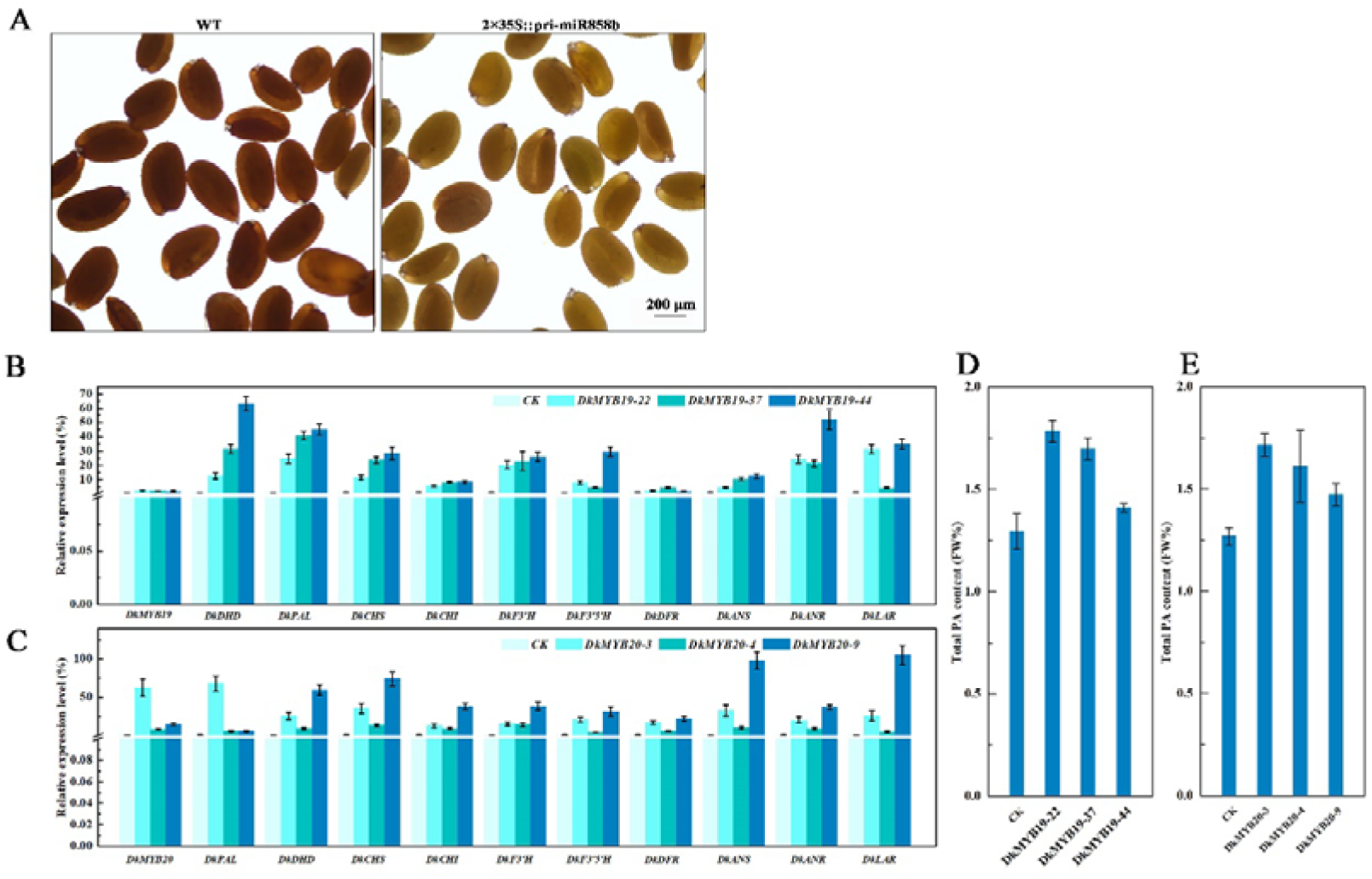
Functional analysis of miRNA858b and *DkMYB19*/*DkMYB20* with a stable transformation system. A, Images of wild-type (WT) Arabidopsis seed, and T1 *Arabidopsis* transgenic lines derived from 2×35S::primiR858 transformation WT background are presented. Scale bar: 200 μm. B, D Analysis of the transcript level of 3 *DkMYB19* genes, PA pathway genes and their PA content variation in the transgenic plant of *DkMYB19*. C, E Analysis of the transcript level of 3 *DkMYB20* genes, PA pathway genes and their PA content variation in the transgenic plant of *DkMYB20*.

To test whether the *DkMYB19*/*DkMYB20* genes can regulate PA synthesis, we infiltrated the *DkMYB19* or *DkMYB20* genes into persimmon leaves. For the *DkMYB19* infiltrated leaves, the expression level of *DkMYB19* increased 4-fold compared with the control (Fig. 6C), while the level of *DkMYB20* increased by approximately 8-fold in the *DkMYB20*-infiltrated leaves compared with the control (Fig. 6D). The qRT-PCR analysis revealed that transient expression of *DkMYB19*/*DkMYB20* induced the up-regulation of *DkDHD*/*SDH*, *DkPAL*, *DkCHS*, *DkCHI*, *DkF3’H*, *DkF3’5’H*, *DkDFR*, *DkDFR*, *DkANS*, *DkANR* and *DkLAR*. All infiltrations stimulated a significant increase in both soluble and insoluble PA content (Fig. 6C, D). DMACA staining visualization of fruit wafers after *DkMYB19* or *DkMYB20* overexpression exhibited distinct darker in both the fruit wafers infiltrated with *DkMYB19*/*DkMYB20* as compared with control (Fig. 7A). Meanwhile, the PA content was increasing correspondingly (Fig. 7B). Thus, we concluded that miRNA858b repressed the expression of *DkMYB19*/*DkMYB20* which involved in regulation of persimmon PA synthesis.

### Genetic transformation in *Arabidopsis* wild type and in persimmon leaf callus

MiRNA858b is a conserved miRNA in the *Arabidopsis thaliana*, we transformed 2×35S::pri-miRNA858b into the *Arabidopsis* wild-type. The seeds of the wild-type (WT) and the 2×35S::pri-miRNA858b lines were germinated on MS medium containing 6% sucrose, then they were transplanted into a culture substrate to generate T1 seeds. The seeds coat of the transgenic lines were paler than their brown wild-type counterparts (Fig. 8A). To test the function of *DkMYB19*/*DkMYB20* with a stable transformation system, persimmon was transformed with CaMV35S-sense *DkMYB19*/*DkMYB20* and a kanamycin-selective marker. We obtained 5 transgenic positive seedlings of *DkMYB19* gene and 15 transgenic positive seedlings of *DkMYB20* gene. Three regenerated positive seedlings transformed by CaMV35S-sense *DkMYB19*, named *DkMYB19-22*, *DkMYB19-37*, *DkMYB19-44* lines, whose expression level of *DkMYB19* upregulated 1.34-fold, 0.95-fold, 0.75-fold (Fig. 8B), showed a gentle accumulation of PA (Fig. 8D), qRT-PCR expression analysis of those three *DkMYB19* trasgenic lines revealed the up-regulation of *DkMYB19* accompanied by a proportional increasion in the expression levels of some structural genes of the PA biosynthetic pathway: *DkDHD*, *DkPAL*, *DkCHS*, *DkCHI*, *DkF3’H*, *DkF3’5’H*, *DkDFR*, *DkANS*, *DkANR* and *DkLAR* (Fig. 8B). The three positive seedlings of *DkMYB20* named *DkMYB20-3*, *DkMYB20-4*, *DkMYB20-9* lines, whose expression level of *DkMYB20* upregulated 61.13-fold, 6.74-fold, 13.80-fold (Fig. 8C), also showed an accumulation of PA (Fig. 8E). The expression of the three *DkMYB20* lines revealed the up-regulation of *DkMYB20* promoted the expression levels of the following PA biosynthetic pathway structural genes of the: *DkDHD*, *DkPAL*, *DkCHS*, *DkCHI*, *DkF3’H*, *DkF3’5’H*, *DkDFR*, *DkANS*, *DkANR* and *DkLAR* (Fig. 8C). All of these results indicated that *DkMYB19/DkMYB20* directly acts as a regulator for the PA pathway genes and controls PA biosynthesis in persimmon.

### The interaction between DkMYB19 and DkPK2

We found DkMYB19 could interact with a DkPK2 protein, that is a pyruvate kinase gene maybe a potential roles in the natural loss of astringency in C-PCNA persimmon via the up-regulation of *DkPDC* and *DkADH* expression during the last developmental stage (Guan *et al*., 2016, 2017). The pGBDT7-DkMYB19 and pGADT7-DkPK2 fusion protein were co-transformed to the Y2H furthermore verified the interactions (Fig. S3).

## Discussion

The persimmon is an ideal material for studying PA metabolism because of it could specifically accumulate PA in its tannin cells, which accumulated a large amount of condensed tannins. The types of “tannin cells” include slender, long, oval, suborbicular, polygon, tip shape, and the surface morphology could be divided into echinate, tuberculiform, concave shape, natural smoothness (Yang *et al*., 2007; Zhang *et al*., 2008). In the fruit, the “tannin cells” mainly exist in the flesh in the form of single dispersed cell or bundles and clumps of cells in the mesocarp, in where they are about two times larger than the similarly shape tannin cells in the exocarp, and the “tannin cells” in the mature fruit are about 3 times larger than that in young fruit. The persimmon PA is the phenolic compounds, which has been associated with biological activities, including inhibition and antiviral, antianaphylaxis, prevention of cardiovascular and cerebrovascular diseases, anti-tumor, promotion of immunity and antioxidant activity (Zhang *et al*., 2008). Morever, the persimmon PA is a kind of natural and eco-friendly adsorbent, it could be used to precious metal adsorption (Xie *et al*., 2016; Fan *et al*., 2019), water purification (Fan *et al*., 2016), soil water retention (Luo *et al*., 2019).

The profile of biosynthesis and metabolism of PA in persimmon is basically clear (Akagi *et al*., 2011). Matsuo and Ito (1978) proposed that PA consists of two types of flavan-3-ols: catechin (C) and gallocatechin (GC), and their gallate ester forms, C-3-*O*-gallate (CG) and GC-3-*O*-gallate (GCG). Gu *et al*. (2008) detected 2, 3-cis epicatechin-3-*O*-gallate (ECG) and 2, 3-*cis* epigallocatechin-3-*O*-gallate (EGCG) units existing in persimmon. Most PA units were suggested to be either EGC or its gallate ester form (EGCG) in all four astringent types, PCNA (pollination constant and nonastringent), PVNA (pollination variant nonastringent), PVA (pollination variant astringent) and PCA (pollination constant astringent), though PA composition was suggested as varying among developmental stages (Akagi *et al*., 2009a, 2010b); EC and its gallate ester form (ECG) were also detected during fruit development. C and GC were demonstrated at an early stage of fruit development, but were barely detectable after the middle stage of fruit development (Akagi *et al*., 2009a).

The PA biosynthesis involved the shikimate pathway, phenylpropanoid pathway, flavonoids - anthocyanins pathway and PA pathway (Akagi *et al*., 2009a, 2011). In persimmon, most genes in both the PA and shikimate pathways including the phenylpropanoid pathway had been identified (Akagi *et al*., 2009a). Two MYB-TFs (*DkMYB2* and *DkMYB4*) were two key regulator that be involved in PA biosynthesis in persimmon (Akagi *et al*., 2009b, 2010a). *DkMYB4* was a predominantly regulator controlled PA biosynthesis in Japanese PCNA persimmon, in which the down-regulation of *DkMYB4* expression during the early stages of fruit growth leads to a substantial down-regulation of PA pathway genes, which results in astringency removal. Moreover, it was suggested that the expression of *DkMYB2* could directly binds to the AC-rich *cis*-motifs known as AC elements in the promoters of the two PA pathway genes, *DkANR* and *DkLAR*, control PA metabolism in persimmon (Akagi *et al*., 2010a). In addition, *DkMYB17* (MH210717.1) that maybe involved in persimmon fruit postharvest deastringency (The paper unpublished). Although a sequence analysis showed that *DkMYB20* shared 93% identity with *DkMYB2*, and *DkMYB19* shared 94% identity with *DkMYB17*, but their function involved in regulating PA biosynthesis is unkown. Our data presented here indicated a novel mechanism integrating translational regulation to control *DkMYB19*/*DkMYB20* expression in PA biosynthesis in persimmon.

At present, a major target of the miRNAs in previously reported is the relatively unknown MYB genes. Among them, miRNA858 could target 36% of MYBs and miRNA828 aimed on 15%, with a TAPIR target prediction score of 6. The miRNA858 was initially functionally identified in *Arabidopsis* (Rajagopalan *et al*., 2006), lately in apple (*Malus domestica*) (Xia *et al*., 2012) and cotton (*Gossypium hirsutum*) (Guan *et al*., 2014). In apple, miRNA858 targets up to 66 MYB factors (Xia *et al*., 2012). As miRNA858 may be involved in numerous biological processes and metabolism pathways, its diverse functions remain to be elucidated. Recently, miRNA858 has been shown to regulate *MYB2* gene homologs that function in *Arabidopsis* trichome and cotton fiber development (Guan *et al*., 2014), and several R2R3 MYB TF genes, including *AtMYB12* (At2g47460), *AtMYB13* (At1g06180), *AtMYB20* (At1g66230), and *AtMYB111* (At5g49330) (Addo-Quaye *et al*., 2008), among that the *MYB12* and *MYB111* are associated with flavonoid metabolism in *Arabidopsis* seed (Stracke *et al*., 2007). The miR828 and miR858 were reported to regulate *VvMYB114* to promote anthocyanin and flavonol accumulation in grapes (Tirumalai *et al*., 2019). Sharma *et al*. (2016) integrated the potential role of light-regulated miRNA858a-*MYB* network in flavonoid biosynthesis and plant growth and development. Wang *et al*. (2016) delineated HY5-miRNA858a-*MYBL2* loop as a cellular mechanism for modulating anthocyanin biosynthesis. Moreover, it had been reported that the blockage of miRNA858 induces anthocyanin accumulation by modulating *SlMYB48-like* transcripts in tomato (Jia *et al*., 2015).

The major targets of miRNA in our analysis were two relatively kown MYBs, named *DkMYB19*/*DkMYB20*, and it’s the first time to screen a persimmom miRNA858b that had a significant differential expression in the two miRNA database at the pivotal time points (15 and 20 WAB) of the PA content reduction in Chinese PCNA persimmon (Luo *et al*., 2015)(Fig. S1). A divergent expression patterns of the miRNA858b that display significant up-regulation expression at 20 WAB and 25 WAB, which caused the down-regulation of the two *MYB* transcription factors. These reports are in agreement with the idea that miRNA858b might target a number of R2R3-MYB genes involved in the secondary metabolite pathway, and even a slight variation of miRNA858b expression could exert obvious influence over product accumulation.

Compared with wild type, transient expression of miRNA858b in the ‘Eshi 1’ leaves *in vivo* or fruit wafers *in vitro* accumulated less PA, and the up-regulated of miRNA858b could suppress the expression of *DkMYB19*/*DkMYB20*, so that resulted in the expression decline of key PA pathway genes (Fig. 6A, 7A; 7B). Transgenic experiment of pri-miRNA858b in *A. thaliana* provided further evidence for its function in PA synthesis (Fig. 8A), while transient expression of STTM858b in the ‘Eshi 1’ leaves *in vivo* or fruit wafers *in vitro* accumulated more PA, and the down-regulated of miRNA858b could promote the expression of *DkMYB19*/*DkMYB20*, so that resulted in the expression rise of key PA pathway genes (Fig. 6A, 7A; 7B), suggesting that miRNA858b is a negative regulator in PA biosynthesis. On the contrary, the transient expression of *DkMYB19/DkMYB20* in the ‘Eshi 1’ leaves *in vivo* or fruit wafers *in vitro* could upregulate the key PA pathway genes (Fig. 6C, D; 7A, 7B), which promote the accumulation of PA (Fig. 6C, D; 7A; 7B). The expression of *DkMYB19*/*DkMYB20* in transgenic plants in tissue culture seedlings of ‘Gongcheng Shuishi’ accumulated more PA than the control, the transcription of key PA pathway genes were significantly increased (Fig. 8B, 8C, 8D, 8E). It meant that miRNA858b acts through repression of *DkMYB19* and *DkMYB20* to involve in regulating PA accumulation. Particularly, *DkMYB19/DkMYB20* maybe regulate the expression of *DkLAR* and *DkANR* specifically, the interaction relationship need further validation through EMSA (Akagi *et al*., 2009b). The regulatory activity of MYB TFs in recognizing and binding DNA with high affinity and specificity has been suggested to involve PPIs and post-translational modifications (PTM; Dubos *et al*., 2010). Importantly, the functional domains of the MYB TF consist of a central transactivation domain, and a C-terminal negative regulatory domain, besides the DNA binding domain (DBD) at the N-terminus. DNA-binding TFs, transcriptional co-activators, and proteins that alter the MYB protein’s activity, e.g., in PTM have been suggested to interact with MYB proteins (Ness, 1999). In this study, the Y2H approach used to screen a fruit cDNA library prepared from the CPCNA, identified complete CDS encoding for PA metabolism protein as potential interacting partners. Our Y2H results suggested that the expression of the *DkPK2* gene and the involvement of PPI may have some relevance in PA metabolism regulation.

Based on our data and earlier studies, we proposed a hypothesis model on the regulation of miRNA858b in the PA biosynthesis pathway in Chinese PCNA persimmon (Fig. 9). The result directly signified that miRNA858b repressed the expression of *DkMYB19*/*DkMYB20*, while *DkMYB19* may contribute to regulate the PA pathway genes to control the PA biosynthesis, and DkMYB20 could interact with DkPK2 to regulate the *ADH* and *PDC* involved in the natural loss of astringency in C-PCNA persimmon. Morever, the introduction of transient transformation in persimmon fruit wafers in *vitro* provide a credible evidence of function of miRNA858b and *DkMYB19*/*DkMYB20*.

**Fig. 9.**
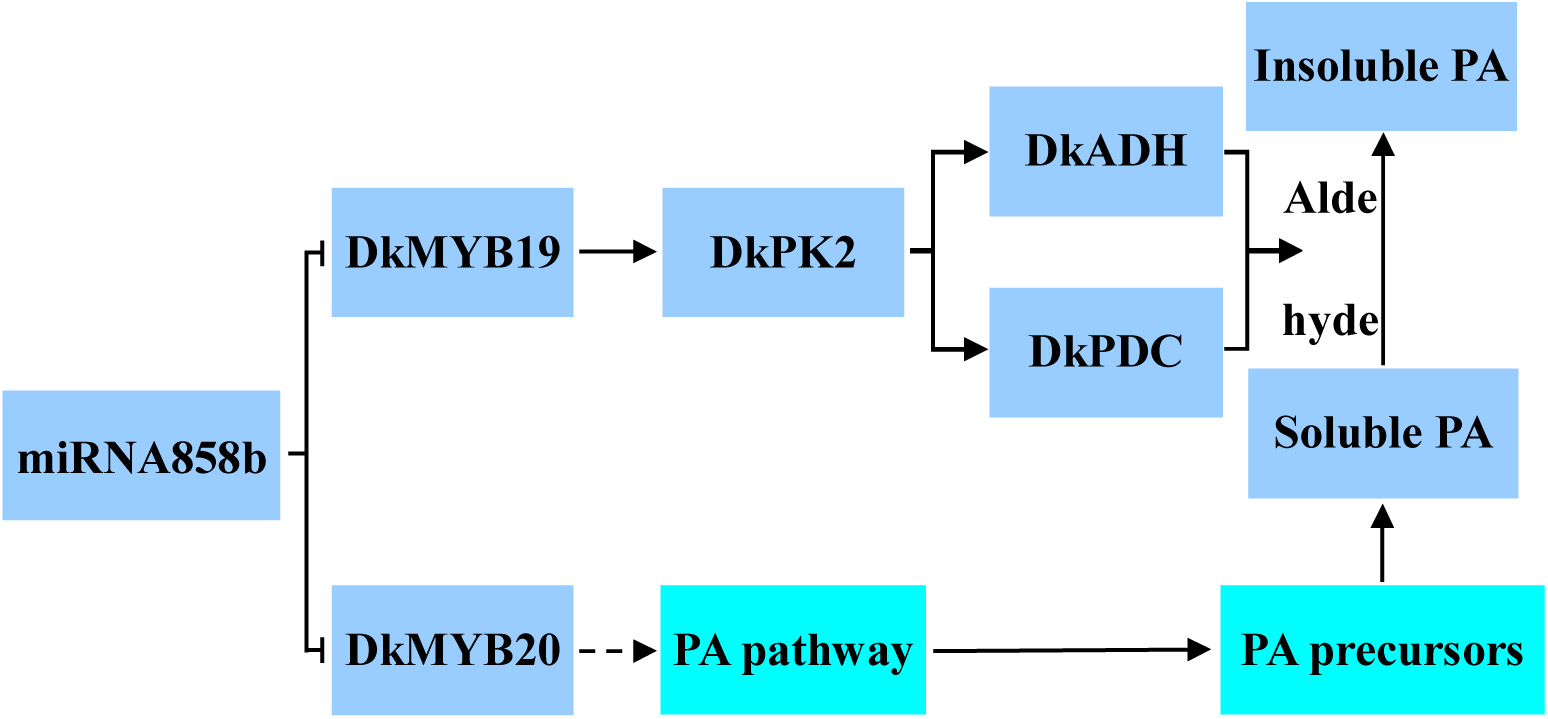
A hypothesis simplified model for miRNA858b-regulated *DkMYB19*/*DkMYB20* repression in PA biosynthesis. In the present study, miRNA858b directly represses *DkMYB19*/*DkMYB20* transcription at the posttranscriptional step. The reduction of *DkMYB19*/*DkMYB20* levels maybe inhibite the expression of the biosynthetic genes of PA precursor genes and key gene in natural loss of astringency in C-PCNA persimmon respectively.

In conclusion, we isolated a miRNA858b and its predicted target genes *DkMYB19/DkMYB20*. miRNA858b interacted with *DkMYB19/DkMYB20* during the fruit development. The function verification demonstrated that miRNA858b-*DkMYB19/DkMYB20* module mediated in the PA accumulation. This study provide genetic evidence for understanding the mechanism of PA metabolism and may contribute for the novel PCNA cultivars breeding.

## Acknowledgements

This work was supported by the National Natural Science Foundation of China (31471846), the Fundamental Research Funds for the Central Universities (2662019PY050) and Hubei Collaborative Innovation Center for the Characteristic Resources Exploitation of Dabie Mountains (2015TD01). The mRNA sequence of *DkMYB19* and *DkMYB20* were submitted to the GeneBank (MN294984 and MN294985).

## Supplementary data

Supplementary data are available at *JXB* online.

Table S1. Primers used in the study.

Fig. S1. qRT-PCR analysis of differentially expressed miRNA during fruit development in ‘Eshi 1’ persimmon. Flesh was collected at 2.5, 5, 10, 15, 20, and 25 WAB. WAB, weeks after bloom. Error bars indicate the standard deviation (n = 3).

Fig. S2. Protein sequence alignment of DkMYB19, DkMYB20 and other R2R3-MYB transcription factors from various plant species.

Fig. S3 Yeast-two-hybrid assay. (A) Self-activation activity detection. pGADT7-T+pGBKT7-53 were used as positive control; pGADT7-T+pGBKT7-lam, pGADT7-T+pGBKT7-lam were used as negative controls; pGBKT7 were used as blank control; (b) Yeast two-hybrid assays of the interactions between DkMYB19 and DkPK2. The empty vectors of pGBKT7 and pGADT7 were used as negative controls.

